# STARTS: A self-adapted spatio-temporal framework for automatic E/MEG source imaging

**DOI:** 10.1101/2024.10.01.616052

**Authors:** Zhao Feng, Cuntai Guan, Yu Sun

## Abstract

To obtain accurate brain source activities, the highly ill-posed source imaging of electro- and magneto-encephalography (E/MEG) requires proficiency in incorporation of biophysiological constraints and signal-processing techniques. Here, we propose a spatio-temporal-constrainted E/MEG source imaging framework-STARTS that can reconstruct the source in a fully automatic way. Specifically, a block-diagonal covariance was adopted to reconstruct the source extents while maintain spatial homogeneity. Temporal basis functions (TBFs) of both sources and noises were estimated and updated in a data-driven fashion to alleviate the influence of noises and further improve source localization accuracy. The performance of the proposed STARTS was quantitatively assessed through a series of simulation experiments, wherein superior results were obtained in comparison with the benchmark ESI algorithms (including LORETA, EBI-Convex, & SI-STBF). Additional validations on epileptic and resting-state EEG data further indicate that the STARTS can produce neurophysiologically plausible results. Moreover, a computationally efficient version of STARTS–smooth STARTS was also introduced with an elementary spatial constraint, which exhibited comparable performance and reduced execution cost. In sum, the proposed STARTS, with its advanced spatio-temporal constraints and self-adapted update operation, provides an effective and efficient approach for E/MEG source imaging.

## I. Introduction

Electrophysiological source imaging (ESI), a technology that could estimate neural electrical activities from noninvasive measurements (i.e., electro- and magnetoencephalography, E/MEG), has attracted growing interests for its capability of detecting brain dynamics at high spatio-temporal resolutionn [1]. In fact, convergent studies have reported promising outcomes achieved by ESI in various research fields (i.e., decoding motor intention [2], localization of epileptic lesions [3] and diagnosis of brain disorder [4]). The distributed model, which represents the sources as current density spread over the entire cortex surface, has been widely used to model the ESI problem [5]. However, solving the distributed model-based ESI is difficult since it is highly ill-posed, i.e., numerous combinations of sources could produce the same E/MEG measurements [6]. Thus, constraints on source properties are required to regularize the solution.

According to biological evidences [7], [8], E/MEG sources are spatially extended and locally homogeneous. That is, the E/MEG are generated by source patches with reasonable extents, wherein the activities in the same patch are highly correlated. Continuous efforts have been made to incorporate such neurophyisiological constraints into source estimation. A widely-used approach is directly applying spatial regularization to obtain the solution with desired spatial properties. Typical example is the low resolution brain electromagnetic tomography (LORETA) [9], which incorporates Laplacian operator to enforce smooth changes between adjacent sources. The major drawback of LORETA is that it tends to over-estimate the spatial extents, resulting in spatially blurred sources. Another approach to obtain source extents involves cortex parcellation techniques. Specifically, the cortex is segmented into several patches prior to source imaging. Then ESI algorithms determine whether each source patch is activated, and reconstruct the source dynamics of the activated patches [10]. The parcellation can be derived from anatomy atlas [11] or data-driven approaches [12]. However, the atlas-based cortex parcellation can hardly reflect individual variations, while the data-driven approaches are often influenced by the noise and the selection of parcellation-related parameters. Advanced constraints to further improve the reconstruction of E/MEG sources are therefore needed to account the complexity and variation of source spatial properties.

Studies of ESI have showed improved performance of source estimations through incorporating temporal information. For instance, autoregressive (AR) model is introduced to account for temporal smoothness of source time courses, which has achieved better performance compared with those only use spatial constraints [13], [14]. The major limitation of these methods is the computational cost due to high dimension of source space. Besides, the mixed-norm framework applies spatial and temporal regularization simultaneously to enforce spatial sparsity and temporal continuity of the solution [15], [16]. Nevertheless, these constraints overemphasize the temporal continuity and may fail to provide additional information for regularizing source activities. Another approach is the incorporation of temporal basis functions (TBFs), where the source can be modeled as a linear combination of TBFs [17], [12], [18]. The TBFs can be either predefined [19] or estimated in a data-driven fashion [12], [17]. Of note, careful selection of TBFs is crucial for accurately characterizing the temporal dynamics and avoiding potential bias of estimated source. In many cases, a residual term is included to capture the source activities that failed to be modeled by fixed TBFs [17]. Alternatively, TBFs can be updated throughout the source reconstruction process [12].

Heuristically, the E/MEG data are contaminated with various noise originated from, e.g., eye motion and muscle movements. In fact, reducing the influence of noises on the reconstructed sources remains barrier to the implementation of ESI even with the help of denoising pipeline. Conventionally, baseline activities are needed to infer noise information. For example, the variational Bayesian factor analysis [20] has been applied to estimate the statistics of the noise from pre-stimulated data for ERP analysis. However, the baseline activities are not available or reliable in many scenarios, such as the analysis of resting-state or single-trial data. To this end, recent work adopt ESI algorithms based on empirical Bayesian learning to estimate the noise in the absence of baseline datas [21], [22]. However, those algorithms only focus on the spatially-constrained model, while the automatic estimation of noise without prior knowledge of baseline data under spatiotemporally constrained ESI framework has not been explored.

Taking into account all the above, we proposed a novel source imaging framework —STARTS to comprehensively model the spatio-temporal properties of E/MEG sources in an automatic data-driven manner. Specifically, we adopt blockdiagonal spatial constraints that models correlation between each dipole and its first-order neighbors to promote local homogeneity of sources. The blocks are automatically determined from data to reconstruct arbitrary source extents. The TBFs of sources and noises, which are updated throughout the reconstruction process, are incorporated to further improve the ESI performance. The performance of the proposed STARTS has been quantitatively evaluated with numerous simulation experiments and two realistic datasets (i.e., epileptic and resting-state EEG). A computationally efficient version of STARTS—smooth STARTS is also introduced. Our findings demonstrate the superior performance of STARTS, underscoring its potential as a powerful tool for revealing brain dynamics in various neuroimaging applications.

## II. Methods

### A. Probabilistic generative model

The scalp E/MEG can be modeled as a linear combination of source activities and measurement noise:

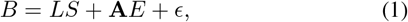

where 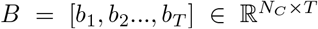 denotes the E/MEG signal, 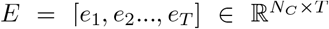 is the noise series, 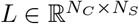 is the known lead-field matrix and 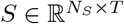 is the unknown sources. *N*_*C*_ is the number of sensors, *N*_*S*_ is the number of candidate sources (*N*_*S*_ ≫ *N*_*C*_) and *T* is the snapshot of E/MEG. 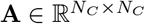 is the mixture matrix for noise that can be estimated if sufficient information about noise source is known. In this paper, **A** is set to be identity matrix *I* since we assume no prior noise information is available. The *ϵ* is a small residual term that satisfies 𝒩 (0, *diag*(*β*)) (𝒩 denotes Gaussian distribution). Here, we represent the source *S* and noise series *E* as a linear combination of TBFs:

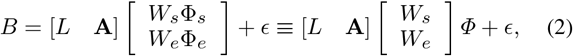

where *Φ* = [*ϕ*_1_, …, *ϕ*_*T*_] ∈ ℝ^*K*×*T*^ is the set of TBFs containing temporal information of both source and noise series, and *K* is the number of TBFs, *W*_*s*_ and *W*_*e*_ are the weights of source and noise TBFs respectively. The identity matrix is used to model the prior covariance of *W*_*e*_. Similarly, prior covariance 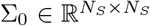 can be used to incorporate spatial constraints into 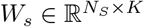 for source estimation.

In Fig. 1(a), we show the graphic representation of the proposed STARTS. Here, a block-manner Σ_0_ is adopted to reconstruct the spatial properties, and the intra-block correlation is learned from data. We model the blocks to capture correlation between each dipole and its first-order neighbors. Such blocks are overlapped, and they can be combined to reconstruct arbitrary source extents (Fig. 1(b)). However, it is hard to incorporate the overlapped blocks directly into source imaging solution. Instead, we transfer the original blocks into non-overlapped covariance matrix [23]:

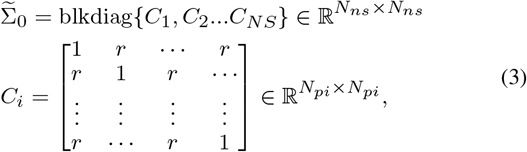

where for each block *C*_*i*_, *N*_*pi*_ = 1+ (the number of neighbors of dipole *i*) and 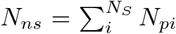. The diagonal elements of *C*_*i*_ are 1 and others are *r* (the intra-block correlation). The 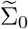 indicates that *W*_*s*_ should be decomposed as:

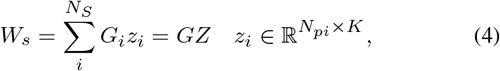

where 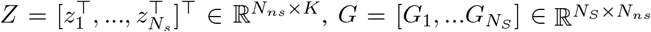, *G*_*i*_ is a zero matrix except that the part from *i*^*th*^ row to (*i* + *N*_*pi*_ − 1)^*th*^ row is replaced by identity matrix, *z*_*i*_ ∼ 𝒩 (0, *C*_*i*_). Substituting Eq. 4 into Eq. 2, we obtain the source model:

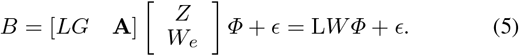

**Fig. 1.**
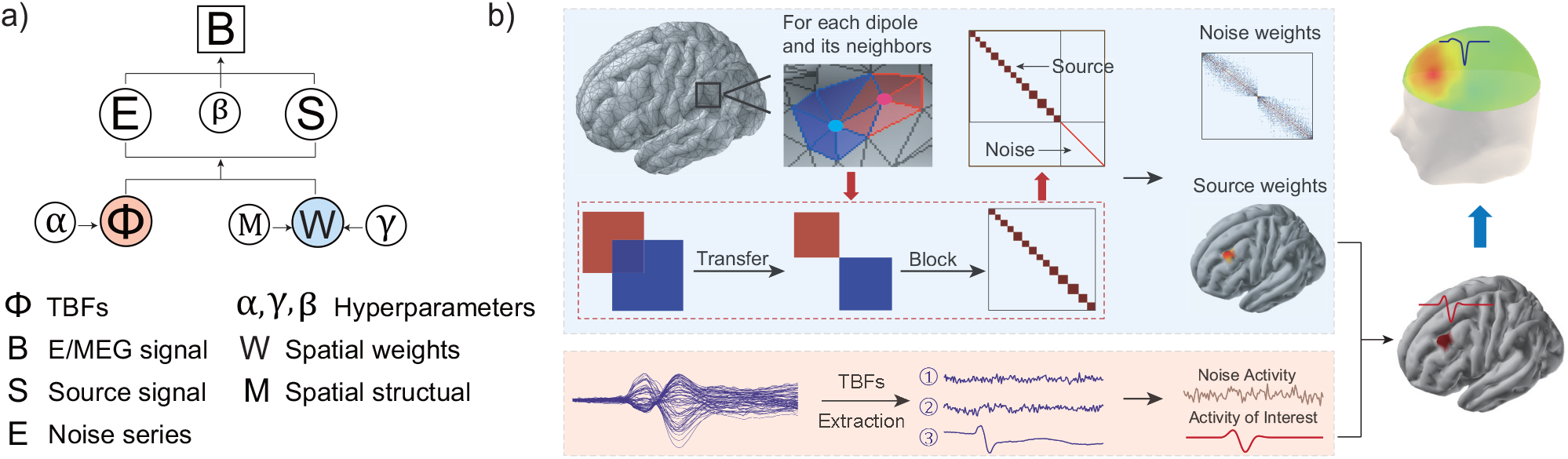
Graphic demonstration of the proposed STARTS. a) Probabilistic model of STARTS. b) Illustration of spatial and temporal constraints used in STARTS. The block-diagonal constraints model the local homogeneity between each dipole and its first-order neighbors. The overlapped blocks are then transformed into non-overlapped ones. TBFs are used to model the time courses of both source and noise activities.

Finally, the source imaging probabilistic model can be expressed as:

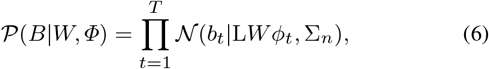

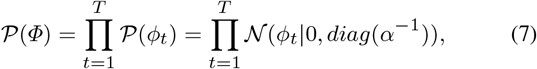

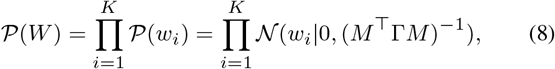

where the augmented prior covariance matrix (*M* ^⊤^Γ*M*)^−1^ is:

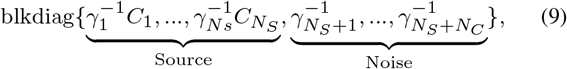

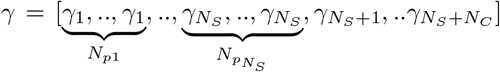 is a preci-sion vector of non-negative hyper-parameters (Γ = diag(*γ*)), *α* = [*α*_1_, …, *α*_*K*_] is a precision vector controlling the relative contribution of each TBFs, *M* is the spatial covariance basis, and 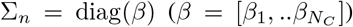) is the prior diagonal matrix of residual term *ϵ*.

### B. Solution to source imaging model

Under the probabilistic model (Eq. 6 Eq. 8), solution to the source imaging problem is the posterior distribution 𝒫 (*W, Φ*|*B*). In this article, we employ the variational Bayesian (VB) to estimate the approximated posterior distributions that match to the true posteriors (i.e., 𝒬 (*W, Φ*) ≈ 𝒫 (*W, Φ*|*B*; *γ, α, β*)), which is calculated by maximizing the free energy [24]:

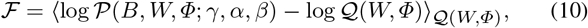

where ⟨·⟩_*q*(*x*)_ = ∫ · *q*(*x*)d*x* denotes expectation with respect to *q*(*x*). We further assume the 𝒬 (*W, Φ*) factorizes over groups of parameters under mean-field approximation:

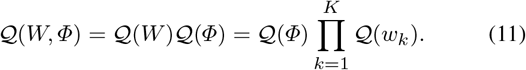

The VB estimates posterior distributions by maximizing free energy ℱ with respect to factorized distributions 𝒬 (*W*), 𝒬 (*Φ*) and hyper-parameters *γ, α* in a expectation-maximization (EM) manner. In the E-step, the posteriors 𝒬 (*W*), 𝒬 (*Φ*) are calculated given the current estimates of hyper-parameters. Then in M-step, the hyper-parameters *γ, α* are updated to maximize the model likelihood (detailed derivations of the update rules are presented in the Appendices). The update of *w*_*k*_ is:

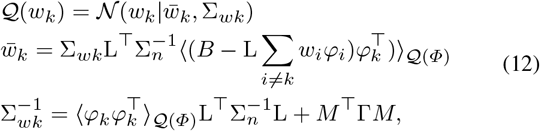

where *φ*_*k*_ is the *k*^*th*^ row of *Φ* and 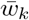 is the posterior mean of *w*_*k*_ (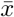 denotes the posterior mean of *x*). The update of *Φ* is:

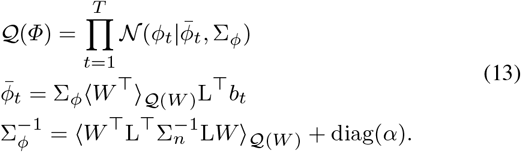

The updates of *γ, α* can be obtained by maximizing ℱ with respect to each hyper-parameter while assuming the posteriors being fixed. Notably, the parts of ℱthat depend on *γ, α* are ⟨𝒫(*W* |*γ*)⟩_𝒬(*W*)_ and ⟨𝒫(*Φ*|*α*)⟩_𝒬(*Φ*)_ respectively, thus we get:

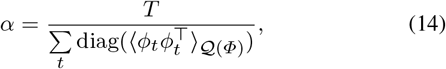

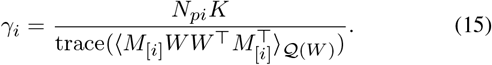

The expectation of quadratic terms in Eq. 12—Eq. 15 is:

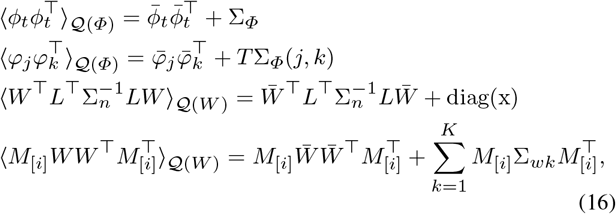

where *x* ∈ ℝ^*K*×1^, 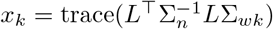, *M*_[*i*]_ is the part of *M* corresponding to the *i*^*th*^ block of (*M* ^⊤^Γ*M*)^−1^ (Eq. 9). In practice, the convergence rate of *γ* is slow with EM-based update rule. To this end, we adopt the convex-based update rules to accelerate convergence [25]. Given the updated posteriors 𝒬 (*W*) and 𝒬 (*Φ*), the ℱ is (details is provided in the Appendices):

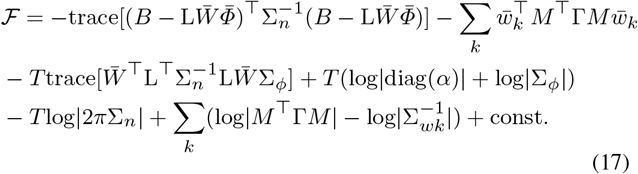

The parts of ℱ depend on *γ* is:

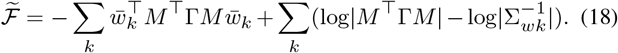

Let 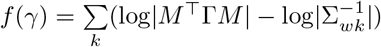, since *f* is convex in *γ*^−1^, it can be expressed as maxima over lower-bounding hyperplanes [25], [18]:

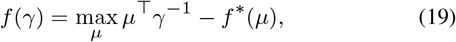

where *f*\* denotes the convex conjugates of 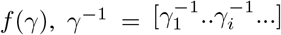, *µ* is the auxiliary variables and is obtained as 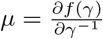. The update of its elements is:

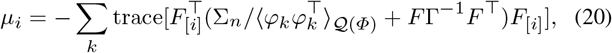

where *F* = L*M* ^−1^, and *F*_[*i*]_ is the part of *F* corresponding to the *i*^*th*^ block of (*M* ^⊤^Γ*M*)^−1^ (Eq. 9). Substituting Eq. 19 into Eq. 18 and setting the gradient 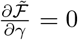, the update of *γ* is:

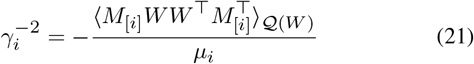

The block structures can also be learned from data by taking the derivation of ℱ with respect to each spatial basis *M* or *C*_*i*_ (which is 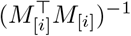) while fixing all other variables. To avoid overfitting and improve the stability of algorithm [23], we enforce all blocks to have the same form (Eq. 3), and to have the same intra-block correlation *r*. Thus, we only need to learn the *r* and update *C*_*i*_ in the form of Eq. 3. We first update each block structure 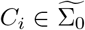 according EM rule (note that the part related to *C*_*i*_ is ⟨𝒫 (*W* |*C*_*i*_)⟩ _𝒬(*W*)_):

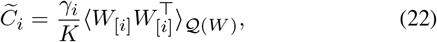

where 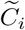 is the updated *C*_*i*_, 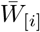 is the part of 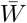 correspond-ing to the *i*^*th*^ block of (*M*^⊤^Γ*M*)^−1^. Then we calculate the averages of the elements along the main diagonal and main sub-diagonal of each 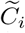, i.e., 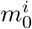 and 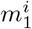, and average for all the blocks to update the *r*:

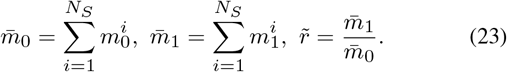

Our primary results, as well as findings from [23] show that the empirical update of *r* guarantees the convergence of algorithm. Finally, the residual parameter *β* is also updated by taking the derivative of Eq. 17 with respect to *β*:

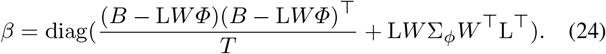

### C. Implementation details

The proposed STARTS can automatically learn the parameters from data, thus it is robust to initialization. For instance, the values of *W* and hyperparameters *α, γ, β* can be simply set to 1. The initial values of *r* is set to 0.99. The TBF set *Φ* is obtained based on the singular value decomposition of *B*. Specifically, given the measurements 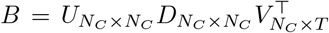, we select the first *K* (e.g., 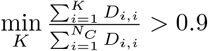) rows of *V* ^⊤^ as the initial TBFs such that *Φ* contains information on both the source and noise activities.

Note that the number of TBFs can be automatically determined by *α*, i.e., the *k*^*th*^ TBF is pruned out if the corresponding 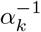 is small: 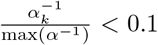. Similarly, we omit the blocks *C*_*i*_ whose relative contribution (measured by 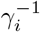) is small: 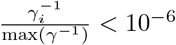.

Notably, update the intra-correlation *r* involves eigenvalue decomposition of 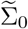 (i.e., 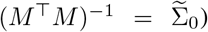 with high dimension of *N*_*ns*_ × *N*_*ns*_ in each iteration, which is computationally cost. Given the form of *C*_*i*_, a faster update can be obtained:

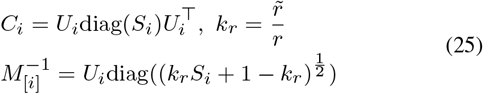

In addition, we can replace the block-diagonal spatial basis *M* with a smooth spatial basis, e.g., surface Laplacian operator *D* − *τ N*, where *D* and *N* are the degree matrix and adjacency matrix of the source dipoles respectively. The parameter *τ* ≈ 1 is set to be 0.99 to ensure the stability of calculation. Then the dimension of *M* is *N*_*S*_ × *N*_*S*_ and the new algorithm (smooth STARTS, sSTARTS) is more computational efficient than STARTS, but at the cost of decreased performance. The sSTARTS works in a similar way as STARTS, except that there is no need to use or update intra-correlation *r*.

## III. Performance evaluation

### A. Simulation protocol

A series of simulation experiments are conducted to evaluate the ESI performance. The simulated data are generated with two source models: event-related spectral perturbations (ERSP) model and neural mass model (NMM). The generation of simulated signals is illustrated in Fig. 2(a). To quantitatively evaluated the performance of ESI algorithms, the metrics are calculated from 100 Monte-Carlo simulations using simulated signals for each condition.

**Fig. 2.**
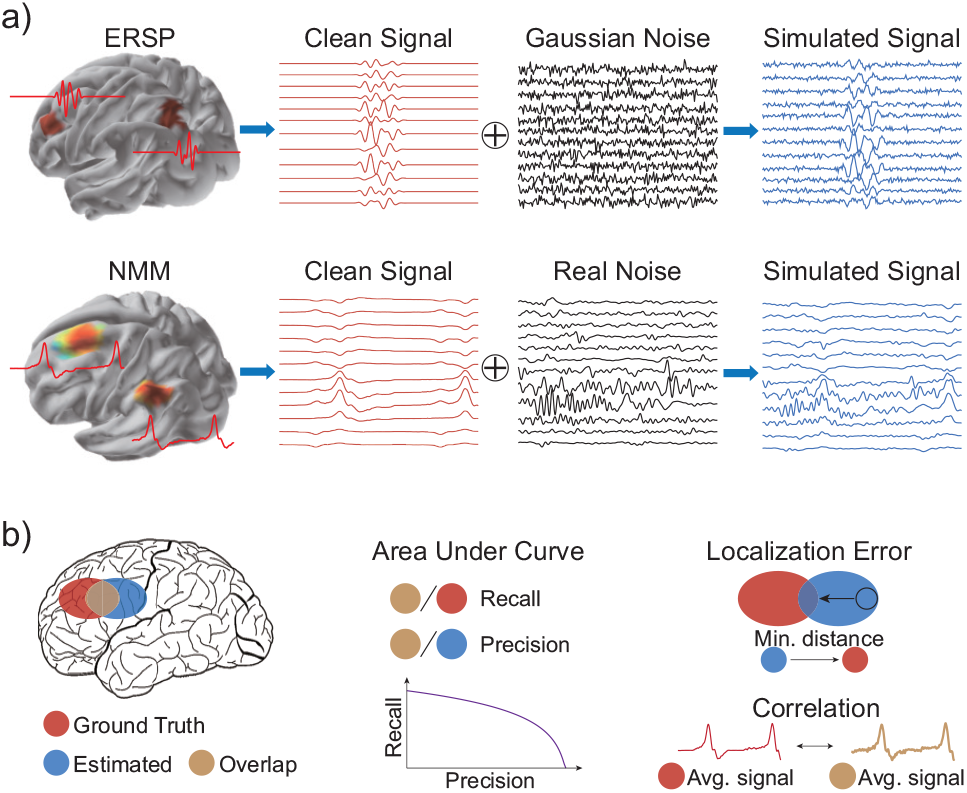
a) The workflow of generating simulated sources with ERSP model and NMM. b) Illustration of performance metrics

#### 1) ERSP

Here, the standard anatomical template — ICBM152 anatomy [26] is used to obtain cortical surface. The cortex is downsampled into 4747 sources and the orientation of each source is restricted to be perpendicular to the cortical surface. The lead-field matrix (53 × 4747) is computed using the Boundary Element Method (BEM) implemented in Open-MEEG (BrainStorm toolbox [27]). For each simulation, the seed voxels are randomly selected. The neighboring dipoles of each seed voxel are subsequently included to generate spatially extended sources. The ERSP source series are generated using SEREEGA toolbox [28]. The simulated EEG are obtained according to Eq. 1, wherein Gaussian noise are added to achieve the desired signal-to-noise ratio (SNR). The SNR is defined as: 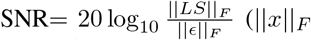 denotes Frobenius norm of *x*). Given that it is often hard to estimate noise activity in real applications, we assume no baseline information is available in this article, and the noise covariance is set to be identity matrix.

#### 2) NMM

To investigate the performance of STARTS under the realistic condition, we generate neurophysiological signals with NMM according to [30]. The template fsaverage5 anatomy is used to get the cortical surface (20484 dipoles), which is then down-sampled to 5000 dipoles. The cortex is segmented into 994 regions with similar size, and we generate simulated sources by setting each region as a seed patch. The spike-like time series of source activity are generated by coupled Jansen-Rit models [31]. Further details on the generation of source activities can be found in [30]. To obtain the simulated EEG, we randomly select seed patches and include the first-layer neighboring dipoles of each patch. The source activities in neighboring dipoles are set to be identical with their seed patches, with a Gaussian weight decay of the magnitude according to the distance of each neighboring dipoles to the center of the seed region. The half maximum of Gaussian weight is set to 50 mm. The EEG data are generated according to Eq. 1, and realistic noise is added to achieve the desired SNR.

### B. Benchmarks and performance metrics

To quantitatively assess the performance of the proposed methods, we select four widely-used validation metrics, including the area under precision-recall curve (AUC) [32], the weighted distance of localization error (wLE), the mean square error (MSE) and correlation (Corr) between the simulated and estimated sources. The AUC measures the sensitivity and specificity of reconstructing source locations and extents. The wLE measures the spatial dispersion and localization error of reconstructed sources. A high value of AUC indicates that the estimated source locations and extensions accurately match to the ground truth. A low value of wLE indicates that the peak values of estimated sources locate inside the truth source patches, and the estimated sources are not spatially blurred. The MSE and Corr assess the temporal error of ESI results. Heuristically, large values of AUC and Corr, along with small values of wLE, MSE indicate good source imaging results. Note that we have scaled the original values of wLE for illustration purpose. The performance metrics is illustrated in Fig. 2(b), and detailed computation of them are presented in the Appendices.

In addition, we compare our proposed methods to three benchmarks, including LORETA, SI-STBF and EBI-Convex.

- LORETA [9]: a widely-used minimum norm based method with spatial smoothness constraint on sources.
- EBI-Convex [22]: an empirical Bayesian framework based solution. It uses augmented lead-field for joint estimation of source and noise, which is solved using the champagne algorithm.
- SI-STBF [12], [33]: a Bayesian method that models source as : *S* = *WΦ*, where the spatial weights *W* and TBFs *Φ* are automatically learned from data. Data-driven cortex parcellation technique [34] is applied to parcel the cortex into several patches, and assumes local homogeneous source activity in each patch.

### C. Real data validation

We also validate the feasibility of the proposed STARTS with two real datasets.

#### 1) Dataset 1

The first dataset contains EEG collection of epileptiform discharges that is public available at ^1^. The 27-channel EEG data are recorded with a sampling rate of 256 Hz. The epileptic spikes are manually marked, and a total number of 58 spikes are averaged for source localization after EEG pre-processing. The individual head model is constructed from pre-surgery MRI, and the cortex is segmented into 6006 dipoles. Detailed information on the datasets and data processing steps can be found at ^1^. The surgical resection outcomes, which serve as the ground truth of seizure generators, can be found at [35].

#### 2) Dataset 2

The second dataset contains resting-state EEG data collected from a 24-year healthy subject (which has been approved by the Institutional Review Board of Zhejiang University). The head model is calculated from individual MRI following the standard pipeline implemented in BrainSuite [36], and the parameters to calculate the lead-field matrix is the same as that used in III.B. The EEG data are recorded from 37-channel wireless system (NeuSen W, Neuracle, China) at a sampling rate of 1000 Hz. The subject first keep eyes open then closed for 5 minutes each. The EEG data are bandpass filtered into 1−40 Hz, denoised by independent component analysis (ICA), rereferenced to common average reference, and down-sampled to 256 Hz. We manually select a 2-second EEG, during which the subject switches from eye-open to eye-closed. We find that the strength of alpha activities at the occipital area are increased as the subject closed eyes. ICA analysis is performed to extract the time courses and spatial weights of those components that are related to the increased oscillation, and the results are used for visual inspection and qualitative comparison.

## V. Results

### A. Simulation results of ERSP model

We first quantitatively evaluate the performance of ESI methods using simulated signals from ERSP model. Fig. 3 shows the reconstructed locations and time courses of five ESI methods. The LORETA yields noisy time courses and it fails to reconstruct the source patch 3. With additional temporal constraints, the sSTARTS produces better performance than the spatially constrained LORETA. The SI-STBF yields blurred results of source patch 1, which may be caused by the erroneous cortex segmentation due to noise. On the other hand, the results of EBI-Convex provide little information on source extents. The performance metrics under different SNR are presented in Fig. 4. The STARTS and sSTARTS could yield precise reconstruction of spatial and temporal source activities even when the SNR is low. The reconstructed sources of EBI-Convex are extremely focal, which is indicated by the low wLE and AUC values. Note that the high MSE of EBI-Convex is also caused by the focal sources, while the high correlation value indicates that the results can provide valuable temporal information. The results demonstrate that the proposed STARTS outperforms benchmark methods, therefore attesting to its robustness and superiority.

**Fig. 3.**
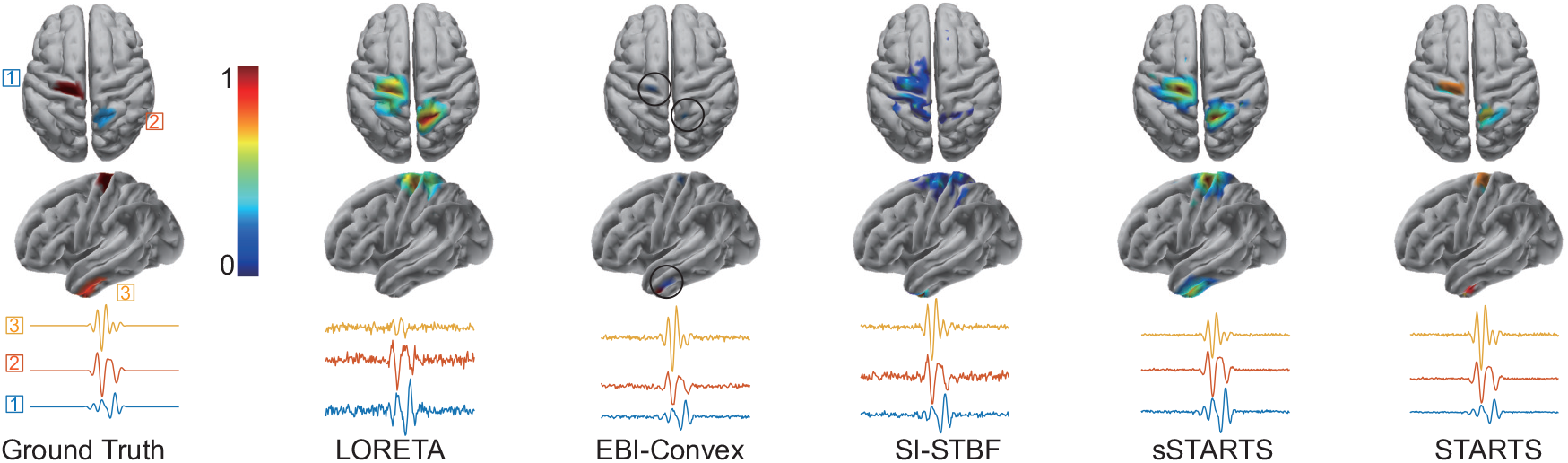
ESI results of ERSP models. The source extents are set to be around 10 cm^**2**^. The source maps are presented as the power of estimated sources (normalized into 0−1). The reconstructed source time series are averaged within each source patch. For LORETA, EBI-Convex, SI-STBF and sSTARTS, the threshold is automatically determined by otsu’ methods [29]. No threshold is used for STARTS. The reconstructed over-focal sources are encircled for better presentation purpose. The SNR is set to be 5 dB.

**Fig. 4.**
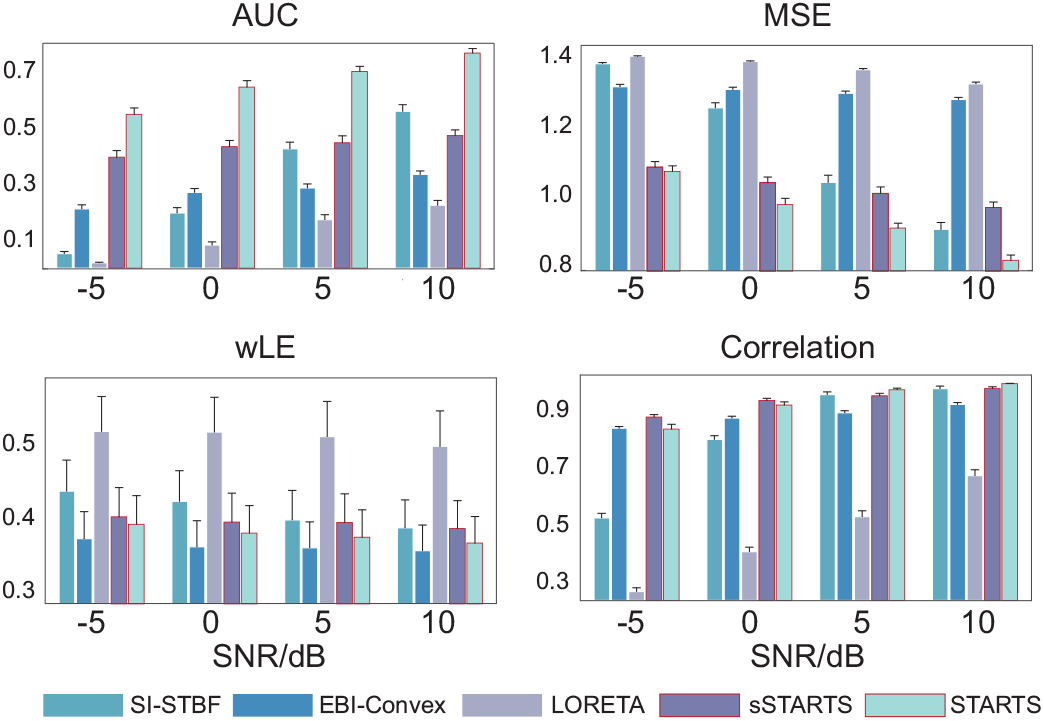
Performance metrics of ERSP source model. The SNR of simulate source ranges from −5−10 dB. The number of source patches is 2. The area of each source is around 10 cm^**2**^. The results are presented as mean **±** SEM (standard error of the mean). The metrics of STARTS and sSTARTS are highlighted.

### B. Simulation results of NMM

We then evaluate the ESI performance using simulated signals from NMM. The NMM is used to generate spike-like sources, and realistic noise is added to generate biologically plausible signals. The mean ± standard derivation of simulated source area is 12 ± 5 cm^2^. Notably, the simulated sources have non-uniform amplitude within each source patch. Fig. 5 presents an example of source imaging results, wherein the STARTS, sSTARTS, and EBI-Convex provide reliable estimation of source locations and time course, while LORETA and SI-STBF show poor performance. Fig. 6 presents the performance metrics under various SNR. The proposed STARTS shows high spatial specificity (indicated by high AUC and low wLE value) and low temporal error (indicated by high correlation and low MSE). The sSTARTS achieve comparable performance with STARTS, while the estimated sources of LORETA and SI-STBF are highly influenced by the noise. The simulated results indicate that the proposed STARTS can reconstruct source activities accurately even if the signals violate the Gaussian assumptions of the algorithm.

**Fig. 5.**
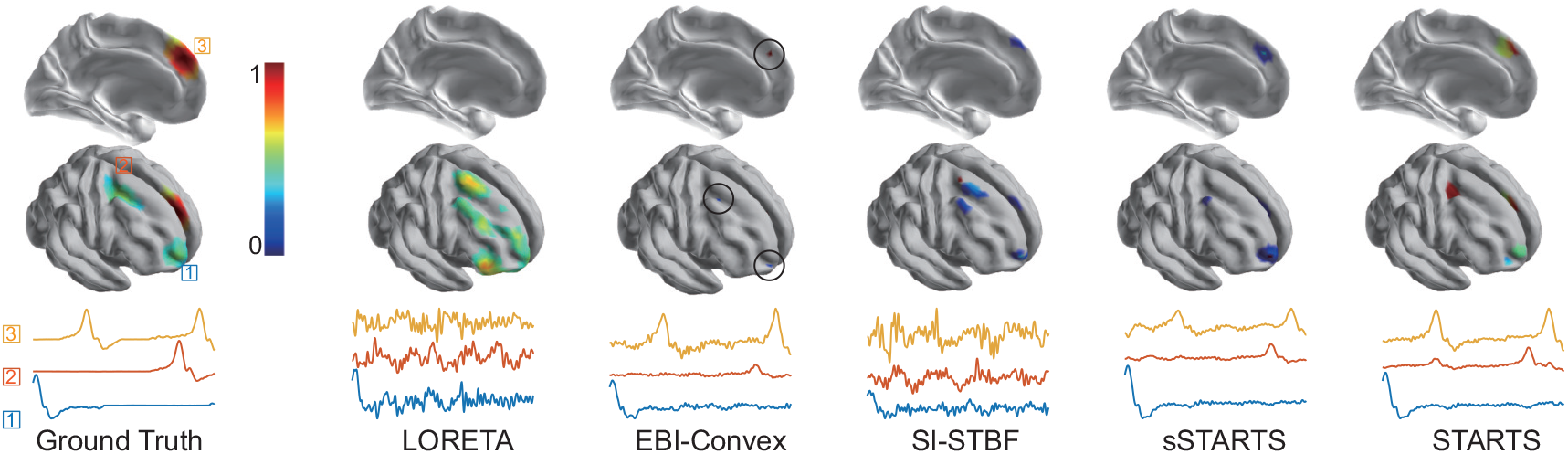
ESI results of NMM. The source extents are 10 −15 cm^**2**^. The source maps are presented as the power of estimated sources (normalized into 0 −1). The reconstructed source time series are averaged within each source patch. For LORETA, EBI-Convex SI-STBF and sSTARTS, the threshold is automatically determined by otsu’ methods. No threshold is used for STARTS. The reconstructed over-focal sources of EBI-Convex are encircled for better presentation purpose. The SNR is set to be 5 dB.

**Fig. 6.**
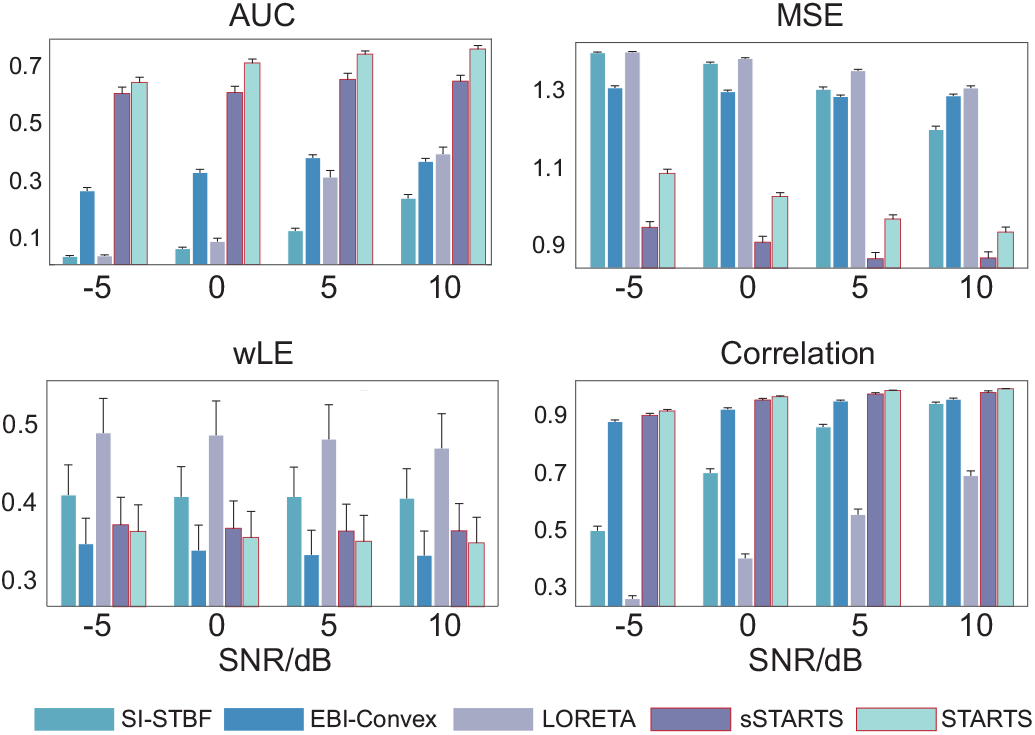
Performance metrics of NMM source model. The SNR of simulate source ranges from −5 −10 dB. The number of source patches is 2. The source areas are around 12 **±** 5 cm^**2**^ (mean **±** standard derivation). The results are presented as mean **±** SEM. The metrics of STARTS and sSTARTS are highlighted.

### C. Results of epilepsy EEG data

The performance of STARTS is further validated with real epilepsy data. Fig. 7(a) shows the topography of average spike following the Brainstorm tutorial. The maximum of spike is recorded at FC1. Fig. 7(b) shows the source imaging results obtained by STARTS along with sSTARTS and EBI-Convex, since the two have showed robust performance in the simulation experiments. The power of reconstructed sources at the maximum of spike (indicated by the blue dash line) is presented. The STARTS localizes two sources: S1 locates at the left frontal lobe and S2 locates at the right occipital lobe, which is consistent with the results obtained by the other two methods. Through investigating the averaged time courses of two sources, we find that S1 is the source related to the epilepsy. Comparing with the results reported in [35], the results of STARTS is in well accordance with clinical outcomes.

**Fig. 7.**
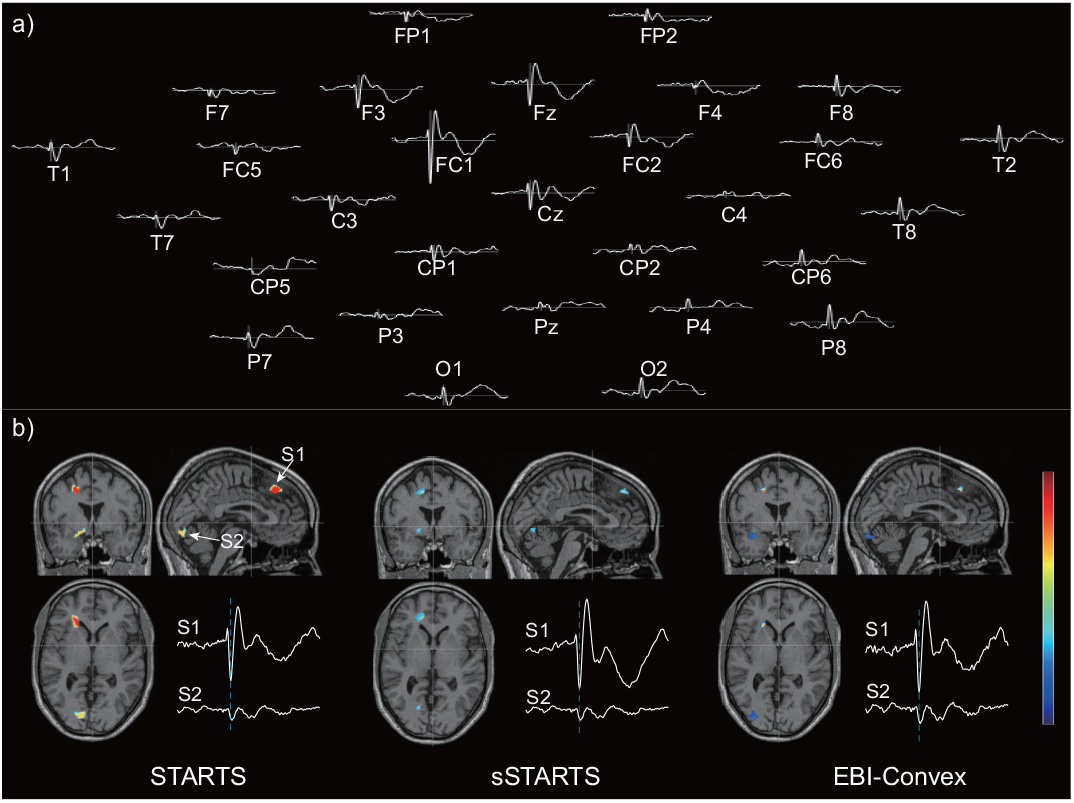
Source reconstruction of epilepsy EEG by three algorithms. a) topography of averaged spike signals. The maximum of spike is recorded at FC1. b) The source imaging result at the peak of amplitude (the blue dash line). The algorithms localize two sources and reconstructs their time courses. The source S1 is related to epilepsy and S2 is irrelevant to epilepsy.

### D. Results of resting-state EEG

In Fig. 8, we present the source imaging results of restingstate EEG by STARTS. The subject first keeps eyes open, and then keeps eyes closed after the time point indicated by the red dash line (Fig. 8(a)). We find a burst of alpha activity after the subject closed eyes. The ICA analysis is performed to find the ICs related to the burst of alpha activity, which is presented in Fig. 8(b). The four ICs account for 72.3% of total power of EEG (IC1:31.7%, IC2:14.4%, IC3:13.4%, IC4:12.8%). Since the ICA decomposition of EEG has been reported to provide valuable information on source activities [37], we use ICA results to qualitatively evaluate the performance of STARTS. The otsu’s method is applied to obtain the threshold to select the dominant source activities. Three source patches and their corresponding time courses are presented in Fig. 8(c). We find that the three sources are relevant to the burst of alpha activity, which account for 67.9% of total power of reconstructed sources. The source 2 and source 3 are located at left/right visual association cortex, and source 1 is located at left primary visual cortex. The results indicate that STARTS produces biologically plausible source estimation of resting-state EEG.

**Fig. 8.**
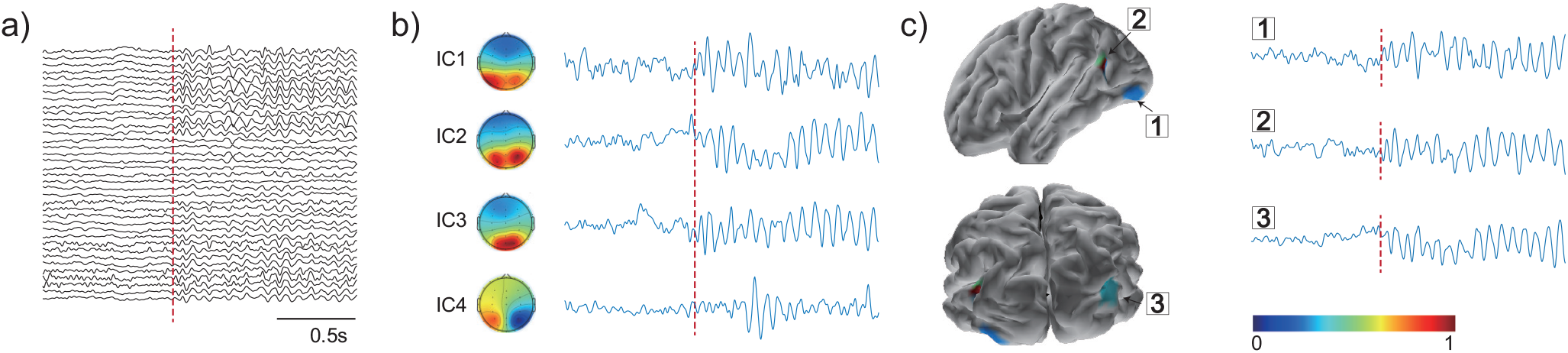
Source reconstruction of resting-state EEG by STARTS. a) The resting-state EEG data. The subject first keeps eyes open, and then keeps eyes closed after the time point indicated by the red dash line. A burst of alpha activity is observed after the subject keeps eyes closed. b) ICA analysis of EEG. The ICs related to the burst of alpha activity. c) reconstructed sources and time courses. The power of sources are presented, and otsu’s method is applied to obtain the threshold.

## V. Discussion

In this article, we propose a fully automatic Bayesian framework STARTS that achieves accurate estimation of extended source in both space and time domain. We incorporate a block-diagonal covariance to facilitate the reconstruction of source extents and correlation of neighboring activities, and include TBFs to improve the accuracy of source localization. Besides, we also integrate the estimation of noise activity into the spatio-temporal source imaging model to enhance its robustness. Simulation results show that STARTS outperforms the state-of-the-art benchmark algorithms. Further validation from realistic data indicates that STARTS can produce biological plausible results. These findings are discussed in detail below. To solve the highly ill-posed ESI problem, STARTS incorporates TBFs in addition to spatial constraints. Consistent with previous work [12], our simulated results (Fig. 4/6) have proved that the inclusion of TBFs helps to improve the source imaging accuracy. We have also provide an additional example to demonstrate the benefits of incorporating TBFs in the appendices (Fig. 9). The ESI results of STARTS, EBI-Convex, and BEBI (an extended version of EBI-Convex algorithm using the same spatial constraint as STARTS) show that the TBFs can facilitate reconstruction of sources, especially the deep ones. Given the apparent benefits of including TBFs in solving ESI problem, practical issues need to be addressed, including how to calculate proper TBFs and to select the number of TBFs. Previous researches have achieved TBFs construction when the signals of interest are available. For instance, Liu et al. used stimulus-evoked factor analysis [38] to estimate TBFs that captures event-related information from baseline signals. In [39], the authors extracted seizure-related activities with ICA decomposition, which were used as TBFs for seizure localization. However, it is challenging to obtain the desired TBFs in most cases. To this end, STARTS adopts a data-driven method to automatically learn and update TBFs from data. Moreover, STARTS can learn the time course of source and noise activity jointly from TBFs, which enhances the robustness of results even in scenarios where the SNR is low. Alternatively, STARTS can utilize user-specific TBFs if the temporal information on sources is available. Our primary results indicate that STARTS still performs well when TBFs contain minimal information on noise (data not shown). In such cases, the estimated noise is negligible, and the values of the noise series are almost zero.

**Fig. 9.**
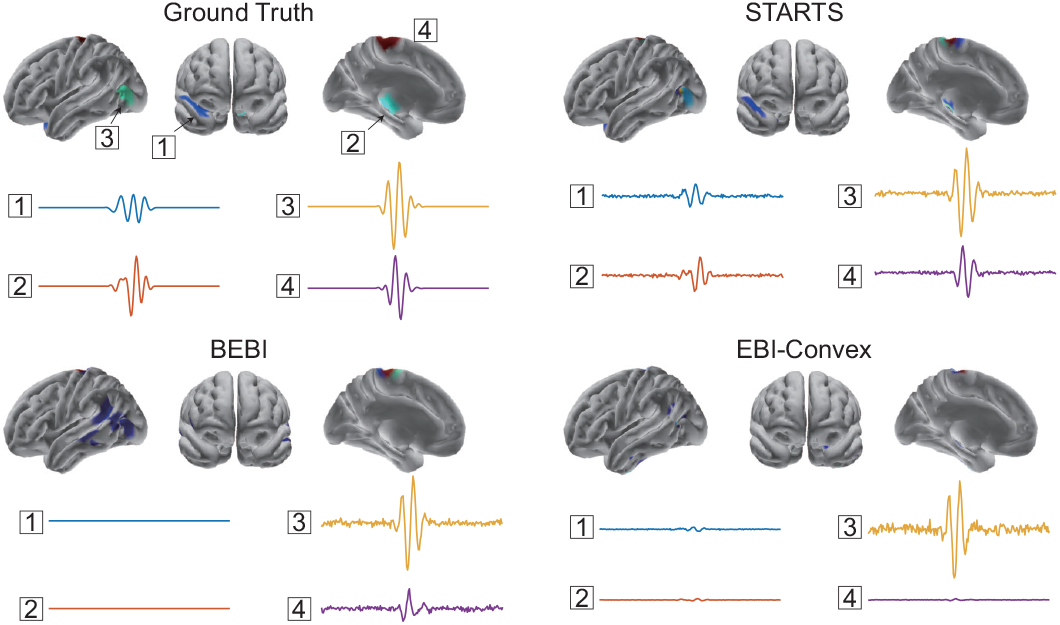
Example of ESI results of STARTS, EBI-Convex and BEBI. The SNR of simulated signals is 5 dB.

Estimation of source extents is another challenge in neuroscience and clinical studies [40]. Spatial constraints, such as the surface Laplacian operator, have been used to enforce smooth changes between adjacent dipoles to obtain extended sources. However, as indicated by the results of LORETA and sSTARTS, the estimated sources tend to be over-smooth, thus a threshold is needed to determine the extents. Another approach, as implemented by SI-STBF, segments the cortex into source patches prior to ESI, and estimates the activity of activated patches [34]. However, the performance of cortex parcellation is sensitive to noise as well as the parameters used to determine the size of each patch. To address these problem, STARTS incorporates block-diagonal constraints to model correlation between each dipole and its first-order adjacent neighbors. This allows correlated blocks to be overlapped, making it possible to reconstruct arbitrary extents. Recently, studies have proved that incorporation sparsity constraint in transformed domain, e.g., edge domain, can achieve accurate estimation of source extents under Bayesian framework [41], [42]. In comparison, STARTS provides an efficient framework to reconstruct source extents in original source domain, which is convenient for inclusion of constraints from other biological findings, such as fMRI results.

The proposed STARTS helps to address practical problems regarding the implementation of ESI, e.g., the estimation of noise. While in most studies baseline signals are required to estimate noise covariance, the proposed STARTS overcomes this limitation by adaptively learning noise activity. Specifically, STARTS employs two strategies to estimate noise. Firstly, STARTS adopts a similar augmented lead-field strategy as [22] to learn the noise time series from TBFs. Secondly, in contrast to [22] where the hyperparameters is fixed to be a small value, STARTS updates the noise/residual covariance in a datadriven manner. Updating the noise covariance is beneficial for the stability of algorithm. After pruning out irrelevant TBFs, the noise covariance are updated to compensate for the missing covariance of the signals. Besides, STARTS can produce clear edges between active and background cortex region without requiring threshold, which is achieved by enforcing most spatial weights of source patches to be zero. Furthermore, STARTS can learn the spatial blocks, intracorrelation, and noise covariance from data, which provides a fully automatic and convenient ESI approach for application. We have also evaluated the performance of a computationally efficient version—sSTARTS. Instead of using block constraint, the sSTARTS incorporates Laplacian operator to ensure spatial smoothness of sources. Our results show that the sSTARTS can achieve comparable performance to STARTS, but a user-specific threshold is required to get the source extents.

There are several aspects that can be further explored to improve the feasibility of STARTS. Firstly, our previous study has indicated that the use of TBFs should consider the non-stationarity of brain activities in practice [18]. The inference of TBFs that account for microstate properties of E/MEG can be further explored in future work. Secondly, instead of assuming noise covariance to be diagonal as in this study, the covariance matrix with complicated structures can be used to improve the estimation of noise [43]. Moreover, STARTS can be extended to incorporate various spatial constraints to improve the accuracy of source estimation. Our results find that the STARTS may under-estimate the source extents when the area of the source patch is large (e.g., 15 *cm*^2^). This is due to the automatic relevance determination in the update of source patches, which tends to yield sparse results. Advanced constraints, e.g., sparse constraint in both source and edge domain, can be incorporated in this framework [39], [44].

## VI Appendix

### A. Derivation of the update rules

In this section we provide the derivation of the 𝒬 (*W*) and 𝒬 (*Φ*). For simplicity, we use *tr*[*a*] to denote the trace of *a*, and 𝒜 to denote *diag*(*α*^−1^). According to Eq. 11-12, the free energy is:

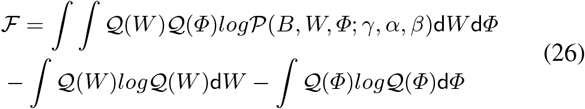

The ℱ with respect to *Φ* can be write as [24], [18]:

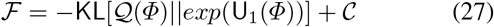

where U_1_(*Φ*) = ∫ 𝒬 (*W*)*log*𝒫 (*B, W, Φ, E*; *γ, α, β*)d*W*, KL[*q*||*p*] denotes the Kullback-Leibler (KL) divergence between *p* and *q*. C = − ∫ 𝒬 (*W*)*log* 𝒬 (*W*)d*W* is a constant term independent of *Φ*. 𝒬 (*Φ*) is updated by setting the KL divergence to 0 [24]:

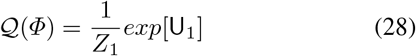

where *Z*_1_ is a normalization constant to ensure the integral of *Φ* is 1. Thus, the update of *Φ* is:

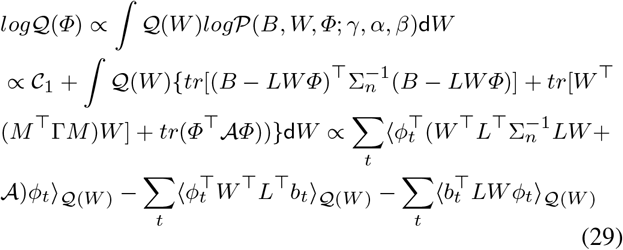

where 𝒞_1_ is independent of *Φ*. The 𝒬 (*ϕ*_*t*_) is Gaussian in *ϕ*_*t*_ as it depends only on *ϕ*_*t*_ and the dependence of *log* 𝒬 (*ϕ*_*t*_) is quadratic. Hence, we obtain the update for 𝒬 (*Φ*) in Eq. 14. Similarly, the update of *w*_*k*_ is:

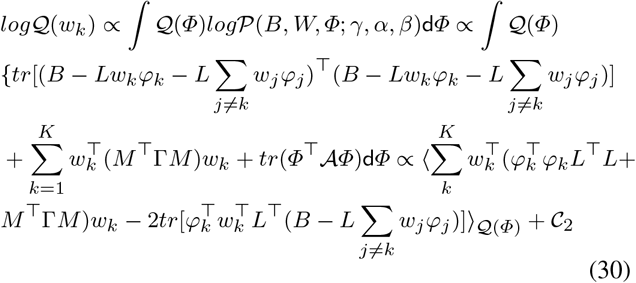

and we obtain Eq. 13.

### B. Derivation of F

Since the 𝒬(*W*), 𝒬(*Φ*) are Gaussian,

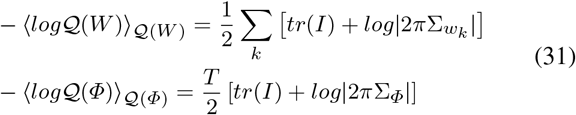

and according to Eq. 7-9, ℱ is:

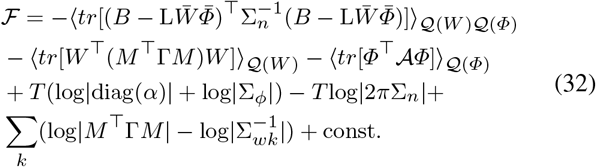

The expectation of quadratic terms in Eq. 33 are:

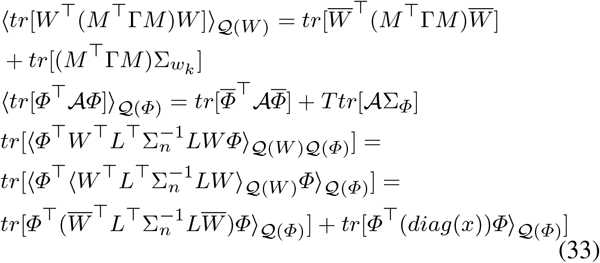

According to Eq. 15, ⟨*tr*[*Φ*^⊤^A*Φ*]⟩_𝒬(*Φ*)_ = *Ttr*[*I*], and according to Eq. 13, 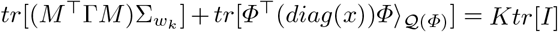. Substituting Eq. 34 into Eq. 33, we obtain Eq. 18.

### C. Calculation of performance metrics

Let *S* and *Ŝ* denote the simulated and estimated sources respectively, *q* = *diag*(*SS*^⊤^) and 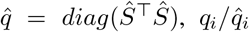 denote the energies of *i*^*th*^ element of simulated/estimated source.

#### 1) area under precision-recall (PR) curve (AUC)

the AUC is the area under PR curve, which is a plot of Recall over the Precision of the estimations as threshold *T*_*h*_ varies. For a specific *T*_*h*_, the dipole is assumed to be active if 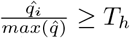. The definition of precision and recall is illustrated in Fig. 2.

#### 2) weighted distance of localization error (wLE)

the localization error (LE) measures the distance of estimated dipoles *i* to the nearest active dipoles of simulated maps (*d*_*i*_, which has been scaled into [0, 1]). The wLE is the sum of the LE weighted by the dipole energies, and we have scaled the wLE for illustration purpose: 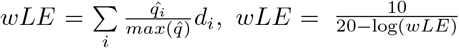

#### 3) mean square error (MSE): 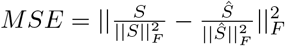

#### 4) correlation (Corr)

the Corr measures correlation (*corr*) of mean time courses in simulated source patches (*J*) of *S* and 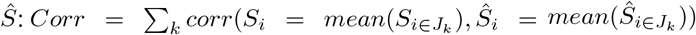 where *J*_*k*_ denotes the *k*^*th*^ simulated source patch.

### D. ESI results with/without TBFs

We provide an additional example here to demonstrate the benefits of incorporating temporal constraints. We show the source imaging results of BEBI, which is an extended version of EBI-Convex algorithm with the same spatial constraints of STARTS. The BEBI aand EBI-Convex both fail to reconstruct the deep sources. In contrast, STARTS successfully find the four sources and reconstruct the time courses.

## Footnotes

1 https://neuroimage.usc.edu/brainstorm/Tutorials/Epilepsy

